# Re-evaluating Reported Pseudolysogeny in Phage T3: T3 and T7 Show Similar Propagation Responses to Nutrient Limitation and Media Switching

**DOI:** 10.64898/2026.08.07.743557

**Authors:** Devyn Del Curto, Brittany Humphrey, Greyson Lasley, Bryce Ricken, Jesse Cahill

## Abstract

Pseudolysogeny is a latent state in which phage development is delayed after infection and has been proposed to promote phage persistence under unfavorable conditions. Virulent phage T3 has been reported to establish pseudolysogeny after infecting starved *E. coli*, then resume lytic replication following transfer to nutrient-rich media, a phenotype linked to the T3 SAMase gene. Here, we revisited the findings of Krueger et al. (1975) to test pseudolysogeny in T3 and examine phage propagation under nutrient-limited conditions. Both T3 and T7 showed impaired propagation under nutrient limitation, with the most stringent conditions causing substantial losses in recoverable infective centers. T3 was modestly more resilient than T7 under these conditions, but we were unable to reproduce the reported phenotype in which T3 remained latent while T7 replicated normally. Supplementation of minimal medium with small amounts of LB supported propagation of both phages, and a repeat experiment designed to more closely match the historical protocol, including post-adsorption reduction of extracellular phage carryover, likewise failed to reveal a T3-specific pseudolysogenic state. Together, our results indicate that, in this experimental system, phage propagation dynamics are more consistently explained by nutrient conditions and media switching than by starvation prior to infection. These findings suggest that the previously reported T3 pseudolysogeny phenotype may depend on additional environmental or methodological factors and underscore the importance of revisiting historically reported phage behaviors using modern controls.

## 1 Introduction

Bacteriophages (phages) are increasingly recognized as effective alternatives to antibiotics for combating drug-resistant bacterial pathogens due to their ability to infect specific target bacteria (Taati Moghadam, Amirmozafari et al. 2020). Unlike antibiotics, phages can selectively target bacterial strains, minimizing collateral damage to the surrounding microbiome (Lin, Koskella et al. 2017, Mu, McDonald et al. 2021, Humphrey, Mackenzie et al. 2023). However, phages that exhibit robust lytic activity in nutrient-rich laboratory conditions often fail to replicate effectively in natural microbiome environments, which are nutrient-limited due to competition for resources (Mitropoulou, Koutsokera et al. 2022, Petrovic Fabijan, Iredell et al. 2023). This discrepancy poses a significant challenge for the development of phage-based therapies. One possible explanation for this phenomenon is pseudolysogeny (PsL), a poorly understood developmental state first described in the 1950s (Fraser 1954) and later studied by Krueger et al. in the 1970s (Krueger, Presber et al. 1975).

Phages exhibit three developmental outcomes upon infecting a host cell. Virulent phages undergo lytic development, replicating and (except for filamentous phages) destroying the host. Temperate phages can either proceed with lytic development or establish lysogeny, a dormant state in which the phage genome usually integrates into the host chromosome and remains repressed (Zhang, Zhang et al. 2022). The lytic/lysogenic “decision”, is processed through a phage-encoded regulatory gene network, which is influenced by factors including multiplicity of infection (MOI, ratio of phage:host cells) and the physiological state of the host (Shao, Trinh et al. 2019). PsL represents a third mode, in which neither lytic development nor “true” lysogeny occurs. Instead, the phage genome remains in a latent/carrier state within the host cell. It is thought that this improves phage persistence under unfavorable conditions until conditions improve to resume lytic replication (Mohammadi and Ely 2025). Despite decades of observations of PsL across diverse phage-host systems, the genetic and molecular determinants of PsL remain poorly described (Abedon 2009). As developed above, the inability to predict phages that exhibit PsL character based on genomic information will likely hinder the effectiveness of phage applications in real-world settings, particularly in nutrient-limited or competitive environments.

It was previously reported that virulent phage T3 establishes PsL upon infecting starved *E. coli* cells but resumes lytic replication when infected cells are transferred to nutrient-rich media (Krueger, Presber et al. 1975). These studies implicated the T3 SAMase gene, which encodes a bifunctional protein with two key activities: overcoming host restriction endonucleases (EcoB and EcoK) and enzymatically degrading S-adenosyl-methionine (SAM), a methyl group donor used in DNA methylation. Krueger’s work suggested that PsL might be regulated by a repressor-like factor responding to methylation changes to the T3 genome (Krüger and Schroeder 1981). In contrast, T7, a closely related phage sharing 89% nucleotide identity over 64% of the genome, lacks the SAMase gene. In nutrient-limited conditions which T3 is reported to show a PsL phenotype, T7 and a SAMase-defective T3 mutant exhibited lytic replication (Krueger, Presber et al. 1975).

Despite the intriguing connections between environmental triggers and epigenetic factors underlying PsL, there have been no follow-up studies building on Krueger’s findings to explore the molecular and genetic basis of PsL using modern tools. By revisiting this phenomenon, we aimed to investigate PsL in greater detail and gain broader insights into its implications for phage biology.

## 2 Materials and Methods

### 2.1 Bacterial Strains and Phages

*Escherichia coli* strain B (ATCC 1130) and bacteriophages T3 (ATCC BAA-1025-B1) and T7 (ATCC BAA-1025-B2) were used for all experiments.

### 2.2 Media

LB broth (Invitrogen #12780-029) was used for bacterial growth and propagation of phage-infected cultures. The broth was prepared according to the manufacturer’s specifications and sterilized by autoclaving. SM buffer was prepared with the following components: 50 mM Tris-HCl, 100 mM NaCl, and 8 mM MgSO₄, with 0.01% gelatin, adjusted to pH 7.5. Where indicated, the buffer was prepared free of gelatin. The buffer was sterilized by autoclaving and used for cell resuspension and phage lysate preparation. M9 minimal media with 0.2% glucose was formulated using the following components per liter: 100 mL of M9 salts, 20 mL of 20% glucose, 10 mL of 0.1 M MgSO₄, 10 mL of 0.01 M CaCl₂, and 860 mL of sterile water. Each component was autoclaved separately to ensure sterility. After autoclaving, the solutions were combined under sterile conditions and mixed thoroughly before use in experiments.

### 2.3 Phage lysate preparation

High-titer lysates of T3 and T7 were prepared by harvesting plaques from 5-10 plates in 5 mL of SM buffer. After centrifugation at 10,000 xg for 10 min, clarified lysates were filtered through a 0.22 µm membrane. Phage titers were normalized to 1 × 10⁹ PFU/mL and residual media was removed by diafiltration using Amicon Filters, 100 kDa MWCO (#UFC9100) and three 15 mL washes in SM buffer. Phage stocks were stored at 4°C and used within one week of preparation.

### 2.4 Preparation of Starved and Non-Starved Cultures

Cultures were grown overnight in LB at 37°C with shaking using a New Brunswick Innova42 incubator at 200 rpm. Subcultures were prepared by diluting overnight cultures 1:100 into fresh LB supplemented with 10 mM MgSO_4_ and 5 mM CaCl_2_ in 20 mL LB in 250 mL baffled flasks. Cultures were grown to an optical density (OD_600_) of 0.6, corresponding to ∼1x10^8^ CFU/mL before use in phage infection experiments. For starvation experiments, *E. coli* cultures were centrifuged at 10,000 × g for 5 minutes, washed twice in sterile gelatin-free SM buffer, and resuspended in gelatin-free SM buffer to a final concentration of ∼7 × 10⁷ CFU/mL. Cells were aerated at 37°C for 1 or 2 hr to induce starvation before phage infection. For non-starved experiments, cells were treated identically with the exception of the starvation treatment.

### 2.5 Phage Infection, Outgrowth, and Quantification

Phage infections were carried out at a multiplicity of infection (MOI) of approximately ∼2 by adding 500 µL of phage stock (1 × 10⁹ PFU/mL) to 4.5 mL of bacterial suspension. The phage-host mixtures were incubated at 37°C for 5 minutes without shaking to facilitate adsorption. For experiments designed to reduce free phage carryover after adsorption, phage-host mixtures were centrifuged at 10,000 × g for 20 sec immediately following the 5-min adsorption step. Approximately 90% of the supernatant was carefully removed without disturbing the pellet, and infected cells were gently resuspended and restored to the original volume in the indicated outgrowth medium. Outgrowth and plaque assay procedures were then performed as described above. This approach was used in a repeat experiment designed to more closely approximate the conditions described by Krueger et al. (1975), in which starved cells were infected after approximately 1 hr of starvation and outgrown in 49:1 M9:LB medium. Following adsorption, the mixtures were diluted to 1 x 10^-2^ by serial dilution followed by a 1:100 dilution in outgrowth media. All dilutions were performed in the respective outgrowth media. For experiments in which phage-host mixtures were transferred from minimal to rich media, a 7.5 mL aliquot was withdrawn using a serological pipette and transferred to a sterile 250 mL baffled flask containing a 2.5 mL of 4x LB. For subsequent timepoints, we monitored the titer of both cultures. At timepoints specified in figures, 250 µL aliquots were collected and further diluted to ensure that ∼30–300 plaques would be countable on each plate. Plaque assays were performed by combining 100 µL of the infective center suspension with 100 µL of overnight culture. The resulting mixtures were added to 1.0% LB top agar maintained at 55°C and immediately poured onto LB plates. Plates were incubated overnight at 28°C to reduce plaque size, facilitating a greater number of countable plaques per plate. Plaques were enumerated the following day. We multiplied plaque counts by 1.333 for plaque assays corresponding to cultures that were transferred to LB to account for the dilution of 7.5 mL into 2.5 mL 4x LB.

### 2.6 Figure Preparation and Statistical Analyses

Graphs were generated using GraphPad Prism 9 software, with raw data initially organized and processed in Microsoft Excel. Statistical analyses reported in Table 1 were performed using Graphpad Uncorrected Fisher’s Least Significant Difference (LSD). For statistical analyses comparing the rate infective center loss for different media or phages, we normalized for slight differences in starting titer between experiments by averaging the titer at time zero and dividing the titers of subsequent timepoints by the average time zero titer before performing the LSD analysis.

**Table 1:** Statistical analyses of select groups of data using Fisher’s LSD (see methods) are reported below along with P-values. Numbers 0, 30, 60, 90, and 120 refer to timepoints (minutes) of sample collection after outgrowth. Summary data shown in GraphPad format: 0.1234 (ns), 0.0332 (*), 0.0021 (**), 0.0002 (***), etc.

| Groups compared | Figure(s) referenced | P-value | Summary* |
| --- | --- | --- | --- |
| T3 LB 0 vs. T7 LB 0 | 1A | 0.9971 | ns |
| T3 LB 30 vs. T7 LB 30 | 1A | 0.551 | ns |
| T3 LB 60 vs. T7 LB 60 | 1A | 0.0507 | ns |
| T3 LB 90 vs. T7 LB 90 | 1A | 0.3971 | ns |
| T3 M9-Sp 30 vs. T3 M9-RO 30 | 1B and 2A | 0.002 | ** |
| T3 M9-Sp 60 vs. T3 M9-RO 60 | 1B and 2A | 0.0376 | * |
| T3 M9-Sp 90 vs. T3 M9-RO 90 | 1B and 2A | 0.1173 | ns |
| T3 M9-Sp 120 vs. T3 M9-RO 120 | 1B and 2A | 0.2261 | ns |
| T3 M9 60 vs. T3 M9 no cells 60 | 1B | 0.0098 | ** |
| T3 M9 90 vs. T3 M9 no cells 90 | 1B | 0.0115 | * |
| T7 M9 60 vs. T7 M9 no cells 60 | 1B | 0.0151 | * |
| T7 M9 90 vs. T7 M9 no cells 90 | 1B | 0.0374 | * |
| T3 M9-RO 90 vs. T3 M9-RO to LB 90 | 1B | 0.1483 | ns |
| T3 M9-RO 120 vs. T3 M9-RO to LB 120 | 1B | 0.292 | ns |
| T7 M9-RO 90 vs. T7 M9-RO to LB 90 | 1B | 0.0118 | * |
| T7 M9-RO 120 vs. T7 M9-RO to LB 120 | 1B | 0.0341 | * |
| T3 M9-Sp 90 vs. T3 M9-Sp to LB 90 | 2A | 0.4176 | ns |
| T3 M9-Sp 120 vs. T3 M9-Sp to LB 120 | 2A | 0.2029 | ns |
| T7 M9-Sp 90 vs. T7 M9-Sp to LB 90 | 2A | 0.1646 | ns |
| T7 M9-Sp 120 vs. T7 M9-Sp to LB 120 | 2A | 0.1185 | ns |
| T3 M9-49:1 90 vs. T3 M9-49:1 to LB 90 | 2B | 0.2763 | ns |
| T3 M9-49:1 120 vs. T3 M9-49:1 to LB 120 | 2B | 0.0712 | ns |
| T7 M9-49:1 90 vs. T7 M9-49:1 to LB 90 | 2B | 0.3039 | ns |
| T7 M9-49:1 120 vs. T7 M9-49:1 to LB 120 | 2B | 0.2378 | ns |
| T3 M9-RO 60 vs. T7 M9-RO 60 | 3A | 0.759 | ns |
| T3 M9-RO 90 vs. T7 M9-RO 90 | 3A | 0.6551 | ns |

## 3 Results

### 3.1 Replication kinetics of T3 and T7 in LB Media

To establish baseline replication behavior, we compared the growth kinetics of bacteriophages T3 and T7 in LB media under non-starved conditions (Fig. 1A). Plaque assays revealed that both phages exhibited nearly identical replication dynamics, with titers increasing from ∼5 × 10³ PFU/mL at time zero to ∼1 × 10⁶ PFU/mL after 90 minutes of outgrowth as observed previously (Hausmann 1973). Intermediate time points at 30 and 60 minutes showed consistent increases in titer, with no significant differences between the two phages (Table 1). These results confirm that T3 and T7 behave similarly in non-starved nutrient-rich conditions in our laboratory, setting a foundation for examining replication rates under starvation conditions.

**Figure 1.**
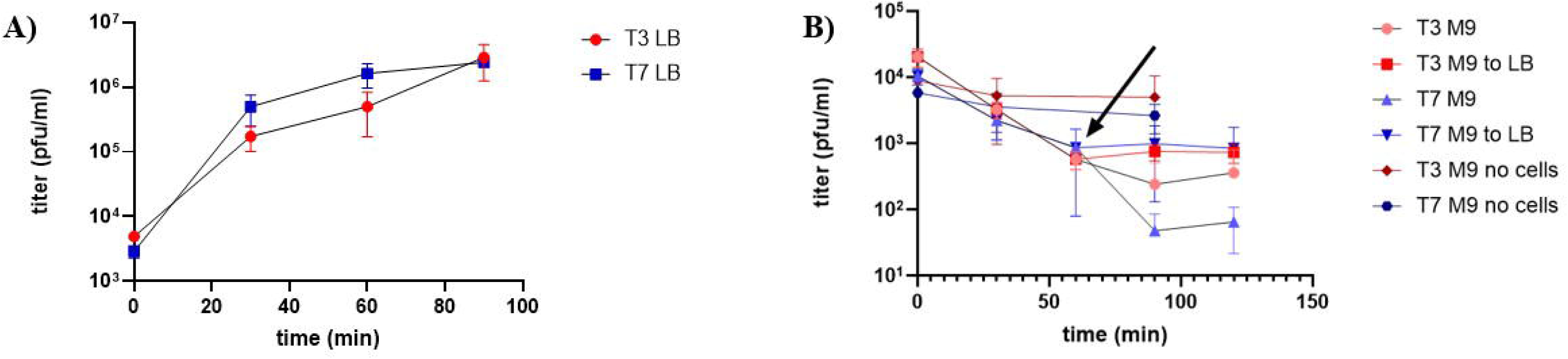
A) E. coli B cells were subcultured to mid-log phase and infected with T7 or T3 at MOI of ∼2. After 5 minutes of adsorption, cells were diluted 10-4 into LB media (time zero). (B) Infections were set up under identical conditions, cells were starved in gelatin-free SM buffer for 2 hours with aeration followed by outgrowth in M9 glucose minimal media. An aliquot of infective centers were shifted to LB media at 60 minutes (denoted M9 to LB and the arrow on the graph) and monitored under identical conditions. Aliquots of infective centers were measured by plaque assay at the time points indicated. Each trace shows the average of 3-4 biological replicates and the error bars represent standard deviation.

### 3.2 Outgrowth in M9 Minimal Media after Starvation Fails to Support Stable Infective Centers for T3 or T7

To investigate the effects of starvation and nutrient-limited outgrowth on phage replication, we compared the behavior of T3 and T7 using *E. coli* cultures starved in buffer prior to infection and subsequently outgrown in M9 minimal media. Both phages exhibited a significant reduction in titer over time, starting at ∼1 × 10⁴ PFU/mL and decreasing by one log after 60 minutes (Fig. 1B). By 90 and 120 minutes, titers dropped further to ∼1 × 10² PFU/mL for T7 and ∼5 × 10² PFU/mL for T3. These results contrast with previous work, which reported that T3 enters a pseudolysogenic state under starvation conditions, maintaining stable titers over several hours (Fraser 1954, Krueger, Presber et al. 1975). In our study, neither T3 nor T7 showed stable infective centers under comparable conditions (i.e., starvation, followed by outgrowth in minimal media), as titers for both phages continued to decline over time. In the aforementioned study (Krueger, Presber et al. 1975), the cells permitted the growth of T7 during starvation, whereas in the current work, T3 and T7 performed identically, with both phages exhibiting log losses in infective centers during outgrowth in M9 minimal media. When aliquots of the phage-host suspensions were transferred to rich media (LB) at the 60-minute timepoint, the rate of titer loss was significantly reduced for T7 after an additional 30 minutes of outgrowth (*P = 0.0118*, Table 1), flatlining at ∼1 × 10³ PFU/mL for both phages at 90 and 120 minutes. This contrasts with previous work, which showed >1 log increase in titer from baseline 30 minutes after infective centers were moved from minimal to rich media.

To determine whether the decrease in infective center recovery in M9 media was due to phage instability—potentially caused by media incompatibility or shearing forces from shaking in baffled flasks—a cell-free outgrowth experiment was conducted (Fig. 1B). Both T3 and T7 showed minimal changes in titer, with significantly higher values at the 60- and 90-minute timepoints in cell-free media (Table 1). These results suggest that the ∼1–2 log reduction in titer is attributable to phage-infected cells rather than the phages themselves.

### 3.3 Nutrient Supplementation to Optimize Conditions for Pseudolysogeny

Given previous observations that trace minerals can significantly improve cell growth in minimal media (Paliy and Gunasekera 2007), we tested whether preparing M9 media with spring water instead of reverse osmosis (RO) water would improve growth outcomes. In this modified media, we observed a small reduction in titer (approximately half a log) at 60 minutes after outgrowth (Fig. 2A), with a modest improvement in recovery of T3 infected cells compared to RO-M9 media (*P = 0.002*, Table 1, T3 M9-Sp vs. T3 M9-RO at 30 minutes). However, while T3 titers remained relatively stable from 60 to 120 minutes, T7 titers decreased by an additional log during this interval. Notably, for T7-infected cells, transferring the cultures to LB media at 60 minutes appeared to improve recovery of infective centers for the next 60-minute interval (Fig. 2A); however, the improvement was not statistically significant (*P = 0.1646* and *P = 0.1185,* Table 1).

**Figure 2.**
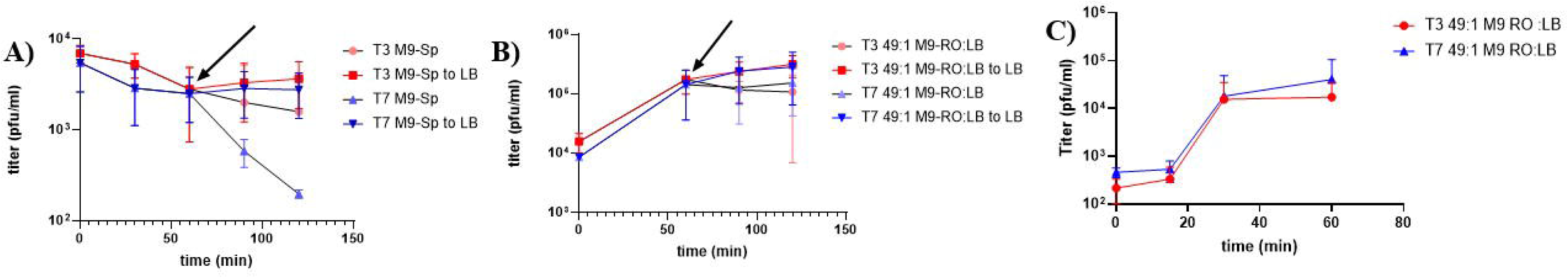
A) E. coli B cells were subcultured to mid-log phase then starved in gelatin-free SM buffer for two hours with aeration prior to infection with T7 or T3 at MOI of ∼2. After 5 minutes of adsorption, cells were diluted 10-4 into M9 glucose minimal media (time zero) made using spring water (M9-Sp). (B) Infections were set up under identical conditions, except outgrowth was performed using M9-RO made supplemented with LB (49:1 v/v).). (C) Infections were set up under identical conditions to (B), except cells were starved for one hour and free phage was removed via centrifugation after the 5 minute adsorption. Panel A and B: An aliquot of infective centers were shifted to LB media at 60 minutes (denoted M9 to LB) and monitored under identical conditions. For all panels, aliquots of infective centers were measured by plaque assay at the time points indicated. Each trace shows the average of three biological replicates and the error bars represent standard deviation.

Building on these findings, we next tested whether supplementing M9 minimal media with small amounts of LB (49:1 v/v dilution ratio M9:LB, a 50x dilution of LB) during outgrowth could further support infective center stability and replication. The goal was to provide sufficient nutrients to support limited cell growth while maintaining the nutrient-limited conditions hypothesized to favor pseudolysogeny. This supplementation stabilized the infective centers and supported replication of both T3 and T7, resulting in a one-log increase in titer after 60 minutes (Fig. 2B). At the 60-minute timepoint, an aliquot of the infective centers was transferred to LB media; however, the titer of T3 and T7 did not change significantly (*P > 0.07* for all comparisons, Table 1) compared to cultures left in M9/LB (49:1) media.

To further test whether free phage carryover after adsorption influenced the observed outgrowth dynamics, we repeated the 49:1 M9:LB experiment under conditions more closely matching the historical report. Starved cells were infected after approximately 1 hr of starvation, and, following the 5-min adsorption period, infected cells were pelleted at 10,000 × g for 20 sec and approximately 90% of the supernatant was removed before gentle resuspension and outgrowth in 49:1 M9:LB media (Fig. 2C). This modification did not reveal a T3-specific pseudolysogenic phenotype. Instead, T3 and T7 exhibited growth patterns qualitatively similar to those observed previously in 49:1 M9:LB medium, with no condition in which T3 remained latent while T7 replicated normally.

While these efforts identified outgrowth conditions that support phage propagation after starvation, including in a repeat experiment incorporating partial removal of extracellular phage after adsorption, we did not observe any condition under which T3 titers remained stable while T7 continued to replicate, as previously reported (Krueger, Presber et al. 1975). However, T7 and T3 did not perform identically in every condition: T7-infected cells consistently experienced greater losses in recoverable titer than T3 in both standard M9 media and M9 media prepared with spring water.

### 3.4 Media-Switching Drives Phage Replication Dynamics in Non-Starved Cells

To further investigate the role of starvation versus media composition in phage replication dynamics, we examined the behavior of T3 and T7 in non-starved *E. coli* cells outgrown in three different media conditions: M9 minimal media prepared with RO water, M9 prepared with spring water, and 49:1 M9:LB mixture prepared with spring water. In RO-M9 media, both T3 and T7 titers showed little change after 60 minutes (Fig. 3A) and moving the infective centers to rich media did not result in a significant change in titer at 90 minutes compared to the cultures in minimal media (analysis not shown). Similar trends were observed for non-starved cells infected with T7 or T3 and transferred to M9-Sp as to M9-RO (Fig. 3B). In contrast, phage-infected non-starved cells outgrown in 49:1 M9:LB media (Fig. 3B) supported T3 and T7 replication, similar to that of starved cells outgrown in 49:1 M9:LB media (Fig. 2B).

**Figure 3.**
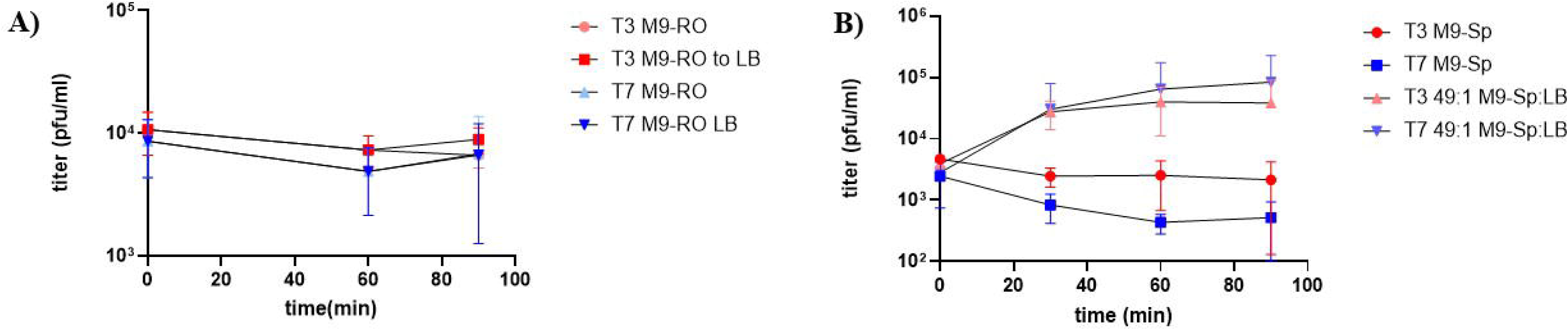
A) E. coli B cells were subcultured to mid-log phase and infected with T7 or T3 at MOI of ∼2 without prior starvation. After 5 minutes of adsorption, cells were diluted 10-4 into M9 glucose minimal media (time zero) (M9-RO). An aliquot of infective centers were shifted to LB media at 60 minutes (denoted M9-RO to LB) and monitored under identical conditions. (B) Infections were set up under identical conditions, except outgrowth was performed in M9-Sp supplemented with LB (49:1 v/v). Each trace shows the average of three biological replicates and the error bars represent standard deviation.

These findings suggest that media-switching, rather than starvation, is the primary factor influencing phage replication dynamics. Both starved and non-starved cells exhibited similar behavior when outgrown in nutrient-limited media, highlighting the critical role of changing nutrient levels for infective center stability and phage propagation. Minor improvements in recovery were observed when spring water was used, suggesting that trace minerals may play a supporting role in phage-host interactions under nutrient-limited conditions.

## 4 Discussion

### 4.1 Revisiting Prior Observations of Pseudolysogeny and Methodological Challenges

Previous studies examined PsL in phage T3 (Fraser 1954), later linking PsL to the SAMase gene and its role in responding to starvation (Krueger, Presber et al. 1975, Krüger and Schroeder 1981). However, these studies lacked detailed methodological descriptions, and the biological components used during that period are no longer available (D. Krüger, personal communication, 10-28-2020). We attempted to replicate these findings using both T3 and T7 under starvation, minimal media, and media-switching conditions. Despite closely following available methods, we were unable to observe a reproducible PsL phenotype in T3. Both phages exhibited similar trends, including loss of infective centers under our most stringent conditions, and in our experimental system phage propagation dynamics were more consistently associated with nutrient switching than with starvation prior to infection. Consequently, this outcome justified our decision not to pursue testing of SAMase mutants, as we could not replicate the critical condition––T7 replication with concurrent T3 latency––reported in prior work (Krueger, Presber et al. 1975).

One notable difference between our study and Krueger’s work is the use of T3-specific antiserum to neutralize free phage and block further adsorption during outgrowth. The use of T7-specific antiserum for T7 experiments as a counterpart to the T3-infected cultures was not reported in the original study. It is possible that T3 antibodies used cross-reacted with T7 (Adams and Wade 1954); however, we observed T7-specific neutralization for a commercially available monoclonal T7 antibody (Merck Millipore #71530, data not shown). Since we lacked a neutralizing T3 antibody, we performed a 10^-4^ dilution after adsorption, as described in the original study into outgrowth media. Dilution is a generally accepted method to reduce further phage adsorption (Hyman and Abedon). It should be noted that the lack of neutralizing antibodies prevented our ability to remove free phage so that our plaque assays were quantifying solely phage-infected cells. However, our data indicate that this is unlikely to be a confounder. We can clearly differentiate conditions in which phage are replicating normally (Fig. 1A, 2B, and 3B) from those that are experiencing a severe reduction in recovery of infective centers (Fig. 1B). We have also identified conditions showing no significant change in phage recovery over time (Fig. 3A). Interestingly, the use of blocking antibodies in the prior work might have introduced a confounding variable on its own. Trace quantities of rich nutrients present in antibody sera would carry over to minimal media during the outgrowth step and might account for the PsL phenotype observed in these studies. Our data indicates “pure” minimal media formulations (in the absence of supplements) do not support replication of phage-infected cells. In other words, antibody sera used in previous work might contribute nutrients to media, similar to what we observed for M9 media supplemented with small quantities of LB (Fig. 2B and Fig. 3B).

Because our initial experiments relied on dilution rather than antibody neutralization to limit further adsorption after infection, we considered whether free-phage carryover might have obscured a T3-specific latent state. To address this possibility, we repeated the experiment using a brief post-adsorption pelleting step (10,000 × g for 20 sec) to reduce extracellular phage carryover before outgrowth, while also more closely matching the approximately 1 hr starvation interval described in the historical study. This additional control produced a result qualitatively identical to our earlier experiments and again failed to reveal a T3-specific pseudolysogenic phenotype (Fig. 2C). Thus, the absence of pseudolysogeny in our study is unlikely to be explained solely by residual free phage after adsorption.

PsL-like behavior for T3 was first described in 1954 by Dorothy Fraser for the Carnegie Institution of Washington Yearbook, who reported that T3-host complexes began to lyse after returning to rich broth at pH 6 but were “more stable” at pH 5 following infection of *E. coli* B starved for 2 hours with aeration (Fraser 1954). However, this work did not use T7 as a non-PsL benchmark and we could not identify growth curve data associated with a PsL phenotype of T3 until a follow-on study that focused on “semitemperate” mutants of T3 (Fraser 1957), which did not examine the previously reported pH effects on the stability of T3-infected cells. Given the sparse methodological detail and uncertainty regarding the genetic background of the T3 mutants, we did not explore pH effects here. However, our approach could be extended to investigate whether pH modulation contributes to phage latency while permitting replication of T7.

### 4.2 Nutrient Levels, Microbiome Competition, and Implications for Phage Therapy

Our findings point to the role of changing nutrient levels for phage replication dynamics, particularly in nutrient-limited environments. In natural microbiomes, competition for resources and fluctuating nutrient availability are common, and these factors could significantly impact the performance of phages sensitive to such changes.

In the most stringent conditions, the recovery of infective centers declined by 1-2 logs over time (Fig. 1B and 2A). Additionally, infective centers were unable to resume lytic development when transferred to rich media (Fig. 3A), as previously reported for the PsL phenotype (Krueger, Presber et al. 1975). However, supplementation with small volumes of LB restored lytic development in both T3 and T7 (Fig. 3B). Together, this suggests that our system is most sensitive to changing nutrient conditions rather than starvation alone.

It would be interesting to determine whether infective center loss is due to the eventual expression of phage-encoded nuclease genes known to degrade the host chromosome ((Krüger and Schroeder 1981). Such findings would raise intriguing questions about the interplay between environmental stress, host cell viability, and the dynamics of phage replication.

Understanding how nutrient shifts and environmental stress influence phage-host interactions is critical for optimizing phage therapy strategies. Natural microbiomes are nutrient-variable, dynamic, and competitive. Phages with PsL phenotypes could result in therapeutic failure unless phages are selected or engineered to be robust in real world settings (Bull, Levin et al. 2019). Future studies could use our methods to sort virulent phages with strictly lytic development from those exhibiting PsL behavior, enabling further investigation of specific phage genes that control the stability of infective centers under nutrient-depleted conditions. Addressing these questions will be essential for engineering resilient phages that are robust and effective in natural microbiome environments. Overall, our results do not support a reproducible T3-specific pseudolysogenic state under the tested conditions and instead indicate that nutrient regime and media transitions are dominant determinants of propagation dynamics in this system. This work highlights the need to revisit historically reported phage phenotypes using modern controls before inferring specific genetic mechanisms.

## 5 Permission to reuse and Copyright

This is an open-access article distributed under the terms of the Creative Commons Attribution License (CC BY). The use, distribution or reproduction in other forums is permitted, provided the original author(s) and the copyright owner(s) are credited and that the original publication in this journal is cited, in accordance with accepted academic practice. No use, distribution or reproduction is permitted which does not comply with these terms.

## 7 Disclaimer

This paper describes objective technical results and analysis. Any subjective views or opinions that might be expressed in the paper do not necessarily represent the views of the U.S. Department of Energy or the United States Government.

## 9 Author Contributions

DDC: Investigation, Data curation, Visualization, Resources

BH: Formal Analysis, Investigation, Writing-review and editing

GL: Investigation, data curation, resources.

BR: Resources, Investigation.

JC: Conceptualization, Funding acquisition, Methodology, Project administration, Resources, Supervision, Validation, Visualization, Writing—original draft, Writing—review and editing

## Acknowledgments

We thank the Cahill Laboratory members and the Sandia National Laboratories’ Environmental Systems Biology and Molecular and Microbiology Department for their valuable input during this study. We thank Chuck Smallwood for reviewing a pre-submission manuscript draft, suggesting edits, and providing technical feedback. SandiaAI Chat, a version of OpenAI’s GPT-4 architecture was used to ideate, structure, inspire, and proofread writing.

## Data Availability Statement

All datasets generated for this study will be made available in supplementary material.

